# Epigenetic and neural correlates of selective social attention across adulthood

**DOI:** 10.1101/2023.07.06.547939

**Authors:** Meghan H. Puglia, Morgan E. Lynch, Madelyn G. Nance, Jessica J. Connelly, James P. Morris

## Abstract

Social isolation is one of the strongest predictors of increased risk of mortality in older adulthood. The ability to form and maintain the social relationships that mitigate this risk is partially regulated by the oxytocinergic system and one’s ability to attend to and process social information. We have previously shown that an epigenetic change to the DNA of the oxytocin receptor gene (*OXTR* methylation) affects the salience of social information in young adults. Little is known about how the oxytocinergic system ages and what effect this aging system has on social cognitive abilities throughout the lifespan. Here we explore age-related differences in the association between neural response during selective social attention and *OXTR* DNA methylation in young and older adults. We find that older adults activate diffuse areas of visual cortex and dorsolateral prefrontal cortex during selective social attention, consistent with the dedifferentiation and compensatory neural activation commonly reported in aging. We find a significant age-by-*OXTR* methylation interaction on neural response when attending to social stimuli in a complex display; young adults display a positive association between *OXTR* methylation and neural activation, replicating our prior finding that young adults with presumed diminished endogenous access to oxytocin recruit regions of the attentional cortex to a greater extent. This association does not hold for older adults. Instead, perceived social support interacts with *OXTR* methylation to influence neural response during selective social attention. These data suggest that environmental factors like social support moderate biological processes in aging and highlight the importance of a lifespan perspective for understanding associations between individual differences in the oxytocinergic system, neural function, and social behavior.

## Introduction

Understanding the neurobiological factors that enable successful social functioning across the lifespan has important psychological and public health implications (Fabiani, 2012). The failure to form adequate social relationships is one of the greatest risk factors for mortality, akin to smoking fifteen cigarettes a day (Holt-Lunstad et al., 2010). Maintaining strong social ties is particularly important for older adults (Matt and Dean, 1993; Seeman, 2000). Older individuals without a strong social support network suffer from an increased risk of dementia, heart disease, stroke, and suicide (Kuiper et al., 2015; Valtorta et al., 2016; Calati et al., 2019). Aging is accompanied by cognitive decline (Salthouse, 2010), especially within the domains of social cognition (Ruffman et al., 2008; Henry et al., 2013) and attentional control (Madden et al., 2005; Zanto and Gazzaley, 2014). The ability to recognize and attend to important social cues including human faces is critical for the formation of social relationships that impact both physical and mental health.

Multiple streams of incoming information simultaneously compete for our attention. Because our perceptual and cognitive systems have a limited processing capacity, we must selectively attend to stimuli that are relevant to the task at hand while simultaneously ignoring distracting information (Desimone and Duncan, 1995). Salient information automatically captures attention without the need to exert volitional or top-down attentional resources. As highly social beings, social stimuli such as faces are often particularly relevant and considered an intrinsically salient class of stimuli (Theeuwes and Van der Stigchel, 2006; Bindemann et al., 2007; Langton et al., 2008; Wang and Adolphs, 2017). The extent to which social information automatically captures attention varies across the lifespan and between individuals.

Our socio-cognitive abilities and motivations change across the lifespan. While affective empathy and social competencies generally improve with age, abilities to detect cues in human motion and accurately attribute mental states to others decrease in older adulthood (Grainger et al., 2019). These changes may be impacted by both age-related motivational changes and changes in attentional control systems. For example, eye-tracking research indicates that older adults attend less to faces during conversations compared to adolescents and young adults (Grainger et al., 2018; De Lillo et al., 2021). According to socioemotional selectivity theory, older adults are more motivated to focus attention on familiar social partners, as compared to younger adults who are more motivated to engage with unfamiliar persons (Carstensen, 1992). However, little research exists examining the neural correlates of changes to social attention across the lifespan and how these changes might impact social support networks.

Individual differences in social attentional abilities across the lifespan may be in part driven by variability within the endogenous oxytocinergic system. Oxytocin is a naturally occurring mammalian hormone and neuromodulator critical for regulating a host of social behaviors across species (Ross and Young, 2009; Carter, 2014). It is hypothesized that oxytocin exerts its effects on social behavior by boosting the salience of social information (Shamay-Tsoory and Abu-Akel, 2016). For example, administration of exogenous oxytocin enhances signal detection for facial expressions (Schulze et al., 2011), biological motion (Kéri and Benedek, 2009), and social vocalizations (Marlin et al., 2015). The actions of oxytocin are dependent upon its receptor, which is expressed at varying levels across individuals. DNA methylation of the oxytocin receptor gene (*OXTR*) is partially responsible for this variability in expression. Our group has identified a cytosine-phosphate-guanine (CpG) site within the promoter of *OXTR*, site -934 (Gregory et al., 2009) (hg38_chr3:8,769,121-8,769,122), that displays wide variability in methylation levels across individuals (Puglia et al., 2015, 2018). Methylation at this site is significantly and negatively associated with *OXTR* expression in the human cortex (Gregory et al., 2009) and is therefore thought to play a regulatory role in gene transcription and the endogenous availability of oxytocin.

We have previously shown in a sample of healthy young adults that differences in neural response during selective social attention is associated with *OXTR* methylation at site -934 (Puglia et al., 2018). Specifically, we found that young adults with higher levels of *OXTR* methylation – i.e., presumed decreased ability to use endogenous oxytocin – recruited regions of the attentional control network to a greater extent when selectively attending to social stimuli within a complex display. The attentional control network is involved in the endogenous, volitional direction of attention and includes dorsolateral, prefrontal, and parietal cortices as key nodes (Hopfinger et al., 2000; Seeley et al., 2007). These results suggest that individuals with presumed diminished sensitivity to endogenous oxytocin (higher levels of *OXTR* methylation) fail to find social information intrinsically salient and therefore engage additional, compensatory neural mechanisms to attend to social information.

While a few studies have begun to investigate how the endogenous oxytocinergic system ages (Ebner et al., 2013, 2014; Huffmeijer et al., 2013; Sannino et al., 2017), no study to date has examined how aging impacts associations between the oxytocinergic system and neural function during social processes. Here we explore how the aging brain maintains attention to social information presented within a complex display, and whether *OXTR* methylation is associated with social signal detection across adulthood.

## Methods

### Participants

All participants provided written informed consent to participate in these studies which were reviewed and approved by University of Virginia Institutional Review Board for Health Sciences Research. 103 (48 males) younger adults aged 18-31 years (*M*=21.14) and 88 (28 males) older adults aged 58-81 years (*M*=68.27) participated in the present study. To control for potential population stratification related to epigenetic testing, only individuals who self-identify as white and of European descent are included in analyses. Older adults were recruited as a subset from the Virginia Cognitive Aging Project (VCAP), a longitudinal aging study ongoing since 2001. VCAP participants were determined to be cognitively normal (*M* = 29.24, *SD* = 0.93) on the Mini-Mental Status Exam (Folstein et al., 1975), and self-reported to be in very good health (Siedlecki et al., 2008; Likert scale from (1) excellent to (5) poor, *M*=1.88, *SD*=0.83). Data from 54 younger adults were collected previously and reported in Puglia et al., 2018. An additional 49 young adults were recruited as a replication sample and the combined young adult sample was used for age-based comparisons for the current study.

### Epigenotyping

Participants provided 8 mL of blood in either mononuclear cell preparation tubes (BD Biosciences, Franklin Lanes, NJ) for assessment of peripheral blood mononuclear cell (PBMC) methylation (younger adults), or PAXGene Blood DNA Tubes (Qiagen, Valencia, CA) for assessment of whole blood methylation (older adults). An independent tissue comparison sample consisting of 156 adult participants (76 males) aged 16-64 (*M*=21.78) years provided both sample types to determine whether methylation values differ across tissue collection methods. DNA was isolated and subjected to bisulfite treatment which converts non-methylated cytosines to uracil and leaves methylated cytosines unmodified. We then amplified a 116-base pair region of *OXTR* containing CpG site -934 (Gregory et al., 2009) (hg38, chr3: 8,769,121) using polymerase chain reaction, and assessed methylated cytosines via pyrosequencing. Reported epigenotypes are an average of three replicates.

### Selective Social Attention Task

Participants underwent a selective social attention task previously described in Puglia et al. 2018 in which participants were presented with double-exposure images of faces overlaid with houses while undergoing fMRI. In alternating 40-s blocks, participants were instructed to attend to aspects of either the face (6 blocks) or the house (6 blocks) in the stimulus while completing a 1-back task. Each block consisted of 10 images and 4 to 5 “same” hits. Images were presented for 1800 ms with an inter-stimulus interval ranging from 200-2400 ms during which a white crosshair was displayed on a black background at center fixation. Participants responded “same” or “different” via button press while the image was still on the screen. Before entering the scanner, participants completed a practice double-exposure 1-back task to ensure they understood task instructions.

The key manipulation in this task is one of selective attention. Although participants are always simultaneously viewing both faces and houses, attending to the faces enhances face-specific neural responses and activation within face-perception regions (O’Craven et al., 1999; Furey et al., 2006; Herrington et al., 2015; Puglia et al., 2018) to a greater extent as compared to the attend houses conditions. Individuals with possible deficits in the intrinsic salience of social information, including individuals with autism spectrum disorder (Herrington et al., 2015) and those with higher levels of *OXTR* methylation (Puglia et al., 2018) require the recruitment of additional attentional control regions to attend to faces relative to houses. We tested for associations between age and task accuracy with logistic regression models predicting proportion of items correct from age in R (Crawley, 2007; R Core Team, 2020).

### Image acquisition and preprocessing

Imaging acquisition details for the original young adult sample are detailed in Puglia et al., 2018. Scanning for the young adult replication sample and the older adult sample was performed at the University of Virginia on a Siemens 3 Tesla MAGNETOM Prisma high-speed imaging device equipped with a 32-channel head-coil. High-resolution T1-weighted anatomical images were first acquired using Siemens’ magnetization-prepared rapid-acquired gradient echo (MPRAGE) pulse sequence with the following specifications: echo time (TE) = 2.98 ms; repetition time (TR) = 2300 ms; flip angle (FA) = 9°; image matrix = 240 mm × 256 mm; slice thickness = 1 mm; 208 slices. Whole-brain functional images were then acquired using a T2* weighted echo planar imaging (EPI) sequence sensitive to blood oxygenation level dependent (BOLD) contrast with the following specifications: TE = 30 ms; TR = 800 ms; FA = 52°; image matrix = 90 mm x 90 mm; slice thickness = 2.4 mm; slice gap = 2.4 mm; 610 volumes of 60 slices coplanar with the anterior and posterior commissures. Stimuli were presented with the Psychophysics Toolbox (Brainard, 1997) for MATLAB (Mathworks, Natick, MA) using an LCD AVOTEC projector onto a screen located behind the subject’s head and viewed through an integrated head-coil mirror.

Data preprocessing was carried out using the FMRI Expert Analysis Tool (FEAT) Version 6.00, part of the FMRIB Software Library (FSL) (Smith et al., 2004). The following pre-statistics processing was applied: motion correction using MCFLIRT (Jenkinson et al., 2002); slice timing correction using Fourier-space time-series phase-shifting; non-brain removal using BET (Smith, 2002); spatial smoothing using a Gaussian kernel of 5.0 mm full width at half maximum; grand-mean intensity normalization of the entire 4D dataset by a single multiplicative factor; high-pass temporal filtering (Gaussian-weighted least-squares straight line fitting, with sigma = 50.0 s). Additionally, each functional volume was registered to the participant’s high resolution anatomical image, and then to FSL’s standard Montreal Neurologic Institute (MNI 152, T1 2mm) template brain using FSL’s linear registration tool (FLIRT) (Jenkinson et al., 2002). Registration from high resolution structural to standard space was then further refined using FSL’s nonlinear registration, FNIRT (Andersson et al., 2007a, 2007b).

### fMRI analyses

Imaging analysis was conducted using FEAT. At the subject level, time-series statistical analysis was carried out using FSL’s improved linear model (FILM) with local autocorrelation correction (Woolrich et al., 2001). Regressors for each condition (Attend Faces, Attend Houses) were modeled by convolving the time course with a double-gamma hemodynamic response function (HRF), adding a temporal derivative, and applying temporal filtering. An Attend Faces > Attend Houses contrast was computed and the contrast of parameter estimates (COPE) from this analysis for each individual was carried forward to higher-level analysis.

Group-level analyses were conducted at the whole-brain level using FSL’s local analysis of mixed effects (FLAME) stage 1 (Beckmann et al., 2003; Woolrich et al., 2004; Woolrich, 2008) with FSL’s automatic outlier de-weighting algorithm applied, which identifies and de-weights outliers within the fMRI data (Woolrich, 2008). *Z* (Gaussianised T/F) statistic images were thresholded using clusters determined by *Z* > 2.3 (p < 0.01) and a corrected cluster significance threshold of *p* < 0.05 (Worsley, 2001). Clusters that survived correction were registered to subject space and mean *Z*-statistic values were extracted for each participant from these clusters. We then tested for outliers by ensuring that the absolute value of the median standardized residuals of the model was < 3 for each data point. Finally, we tested for data points with undue influence on the model using Cook’s distance (D) > 1 (Cook and Weisberg, 1982) as criteria. The removal of outliers and/or influential points did not appreciably change results. All results are presented with outliers and/or influential points removed.

We first conducted a whole-brain analysis to determine the replicability of the association between *OXTR* methylation and task-specific Attend Faces > Attend Houses activation (Puglia et al., 2018) in our new sample of young adults (*n*=49). Two regressors were included in the model – group mean and mean-centered *OXTR* methylation – and contrasts testing for positive and negative linear associations between *OXTR* methylation and Attend Faces > Attend Houses BOLD activation were computed. One outlier was removed from this analysis. All subsequent analyses considered the full young adult sample (*n*=103).

We next tested for group-level differences in task-specific Attend Faces > Attend Houses activation using a two-sample unpaired t-test with task accuracy included as a nuisance regressor to account for differences in task performance across age groups. Three outliers (2 younger adults) were removed from this analysis.

We next tested for an interaction between age group and *OXTR* methylation to determine whether the slope of the linear association between *OXTR* methylation and Attend Faces > Attend Houses BOLD activation varied as a function of age group. Task accuracy was included as a nuisance regressor in this analysis. Eight outliers (3 younger adults) were removed from this analysis.

Finally, in an exploratory analysis we tested for associations between *OXTR* methylation, social support, and neural activation during selective social attention in the older adult group. Participants in VCAP have been completing the Social Support Questionnaire (Shaw et al., 2007) at each study visit since 2009. This questionnaire assesses embeddedness within one’s social network, as well as the amount of social support enacted, provided, and perceived. Scores can range from 13 to 116, with higher scores indicating greater engagement within a more supportive social network. Two participants recruited for the older adult sample in the present study did not have Social Support Questionnaire data. All other older adult participants completed the questionnaire 1 to 5 times (*M*=2.51) since 2009. Average social support scores ranged from 36 to 98.5. We included the mean-centered measures (*OXTR* methylation, Social Support score) and the interaction terms (*OXTR* methylation x Social Support score) as regressors and Task Accuracy as a nuisance regressor in the model, and computed contrasts testing for linear associations between each regressor and Attend Faces > Attend Houses activation. Two outliers were removed from this analysis.

## Results

### *OXTR* methylation levels do not vary by tissue type or age

We first determined that methylation values do not differ across whole blood and PBMC tissue collection types. In the independent tissue comparison sample, PBMC methylation values averaged 46.72% (*SD*=6.33). Whole blood methylation values averaged 46.45% (*SD*=6.82). Methylation values were significantly correlated across tissue types (*r*=.95, 95% CI=[.93, .96], p < .0001), indicating methylation values in our two imaging samples can be compared.

PBMC methylation for the younger adult sample averaged 47.30% (*SD*=6.06). Whole blood methylation for the older adult sample averaged 47.09% (*SD*=10.31). Mean methylation values did not significantly differ across age groups (*t*(146.82)=0.09, 95% CI=[-2.37, 2.59], *p*=.930), although the variance in methylation values was significantly higher for older adults (*F*(95,102)=3.20, *p*<.001).

### Younger adults perform better than older adults on the selective social attention task

A logistic regression model predicting proportion correct from age group revealed a significant effect of age group on task performance (*B* = 0.43, *p* < .0001) such that younger adults performed significantly better on the task (*M*=63.03%, *SD*=20.72) than older adults (*M*=52.63%, *SD*=15.52). Performance on the attend faces and attend houses conditions was significantly correlated for both younger (*r*=.91, *p*<.001) and older (*r*=.85, *p*<.001) adults.

### The association between *OXTR* methylation and neural response during selective social attention replicates in a young adult sample

We obtained a novel young adult sample (*n*=48) to test for the replicability of the association between *OXTR* methylation and neural response during selective social attention we previously identified in young adults (Puglia et al., 2018). We replicate these previous results and find a significant positive association between *OXTR* methylation and Attend Faces > Attend Houses BOLD response in several regions including left dorsolateral prefrontal cortex and right superior parietal lobule (Figure 1, Table 1).

**Figure 1.**
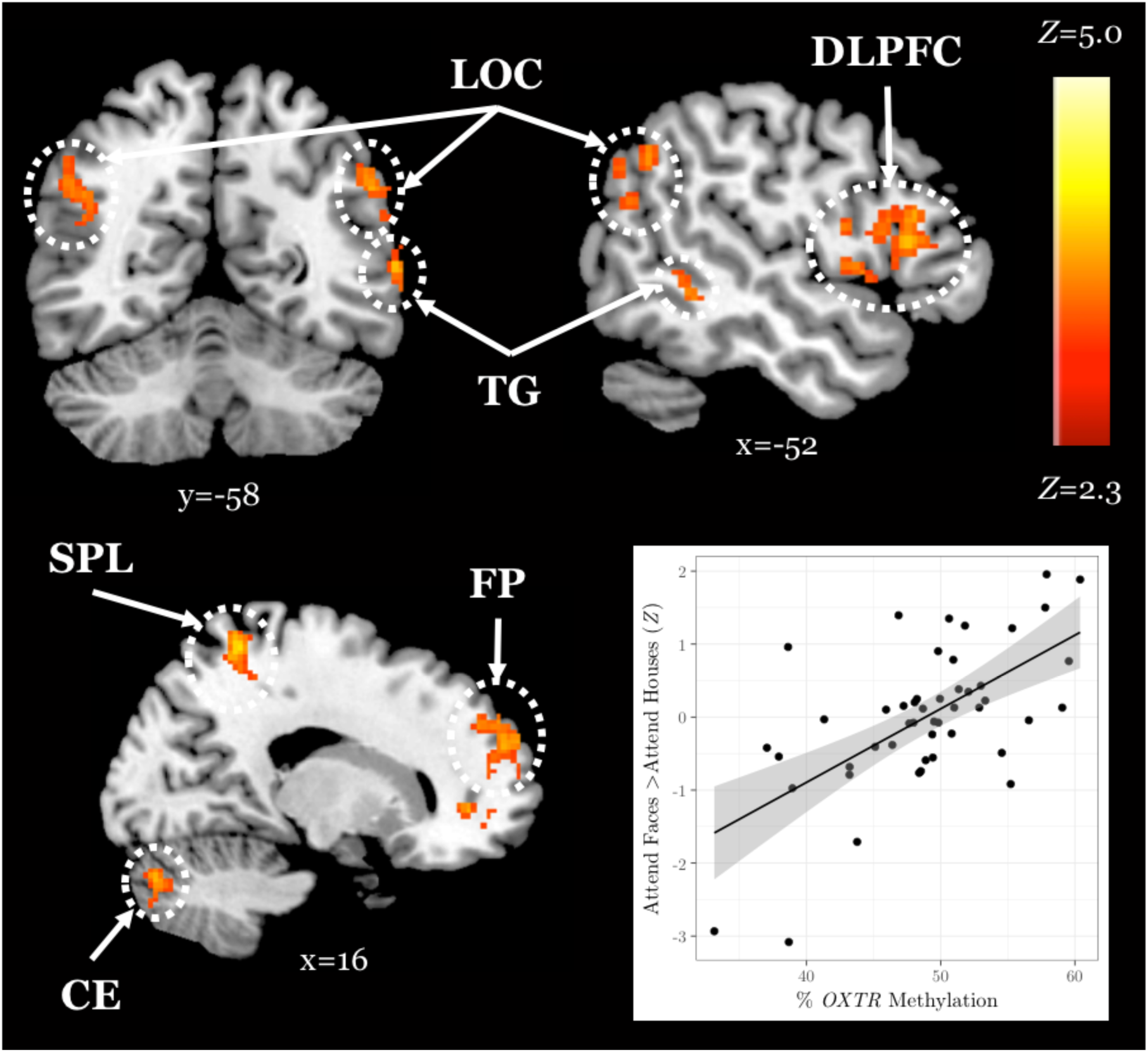
The association between *OXTR* methylation and neural response during selective social attention replicates in a novel young adult sample. *Z* statistic map of regions showing a significant positive association between *OXTR* methylation and Attend Faces > Attend Houses BOLD activity within our young adult replication sample (n=49). Images are whole-brain FWE cluster-corrected at *Z* > 2.3 and depicted in MNI space and radiological orientation. (Inset) Mean *Z* statistic values are plotted against percent *OXTR* methylation for each participant. Solid line depicts the best-fit line; grey shading depicts 95% confidence interval. LOC, lateral occipital cortex; DLPFC, dorsolateral prefrontal cortex; TG, temporal gyrus; SPL, superior parietal lobule; FP, frontal pole; CE, cerebellum.

**Table 1.**
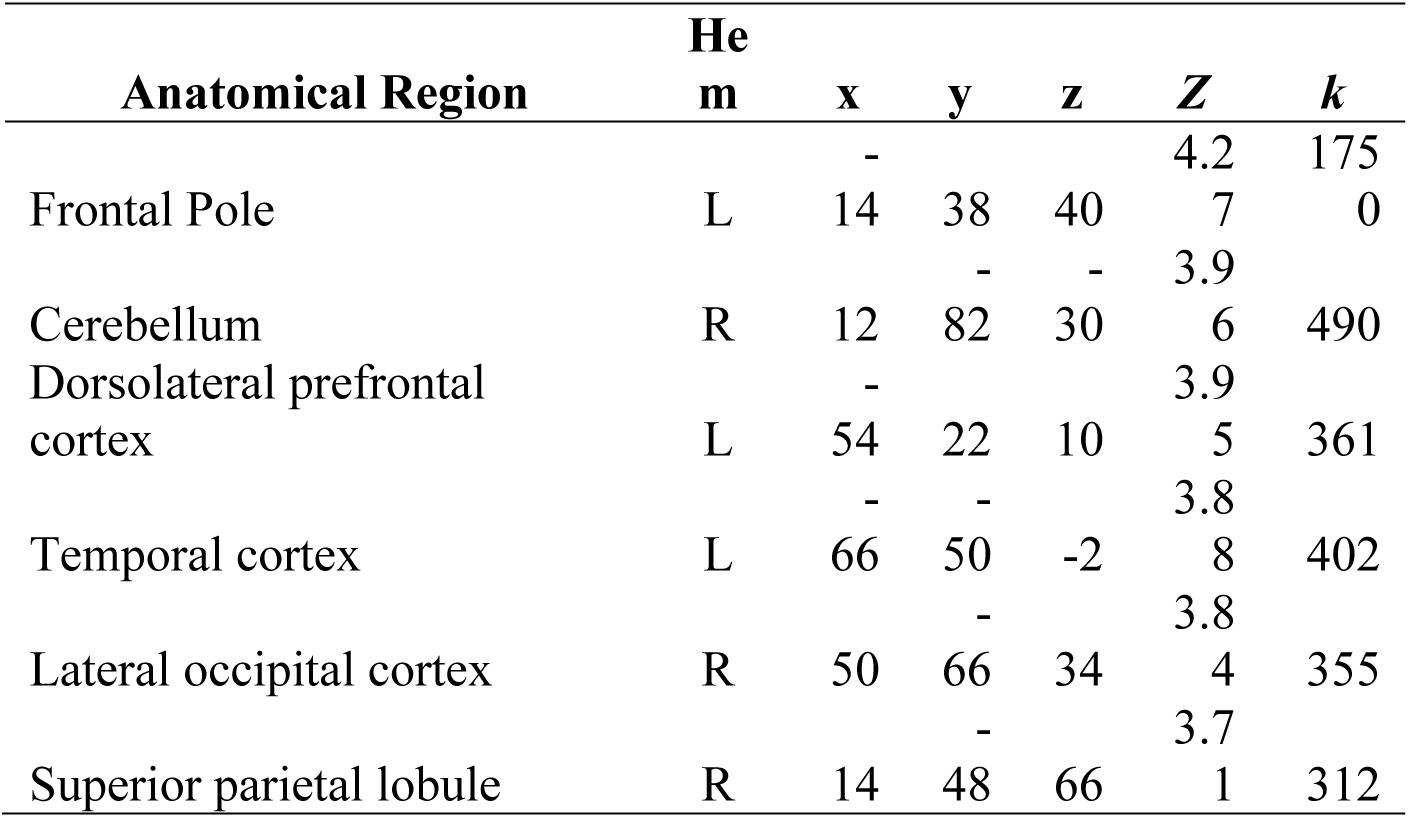
Young adult replication sample local maxima statistics. Hem, hemisphere; L, left; R, right; x, y, z, coordinates of local maxima in MNI space; *Z*, maximum *Z* statistic; *k*, cluster extent.

### Older adults activate a wider area of the brain during selective social attention

When comparing neural response to the selective social perception task across age groups, we find younger adults (*n*=101) activate specific regions of the face perception network including left orbitofrontal cortex and posterior cingulate cortex to a greater extent than older adults (n=87, Figure 2, Table 2). Conversely, older adults show greater activation than younger adults within large regions of visual cortex including bilateral fusiform and lateral occipital cortex (Figure 2, Table 2). Regardless of age, we find participants that performed better on the selective social attention task show greater activation within right dorsolateral prefrontal cortex and right superior parietal lobule (Figure 3, Table 2).

**Figure 2.**
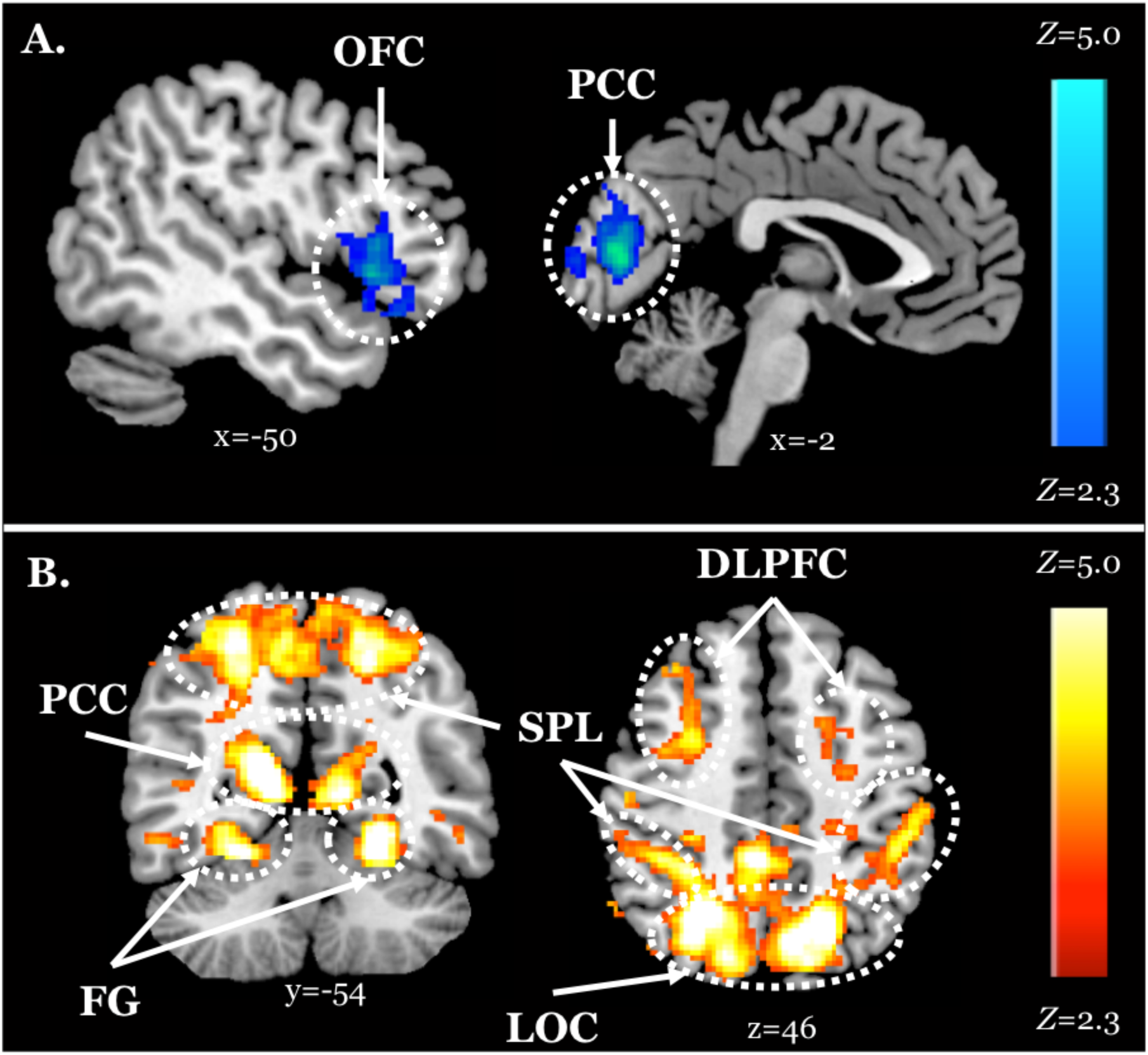
Older adults display neural dedifferentiation during selective social attention. *Z* statistic maps of regions showing significant differences across age groups on Attend Faces > Attend Houses BOLD activity, controlling for task performance. Images are whole-brain FWE cluster-corrected at *Z* > 2.3 and depicted in MNI space and radiological orientation. **A.** Regions that older adults (n=101) activate to a lesser extent than younger adults (n=87). **B.** Regions that older adults activate to a greater extent than younger adults. OFC, orbitofrontal cortex; PCC, posterior cingulate cortex; DLPFC, dorsolateral prefrontal cortex; SPL, superior parietal lobule; FG, fusiform gyrus; LOC, lateral occipital cortex.

**Figure 3.**
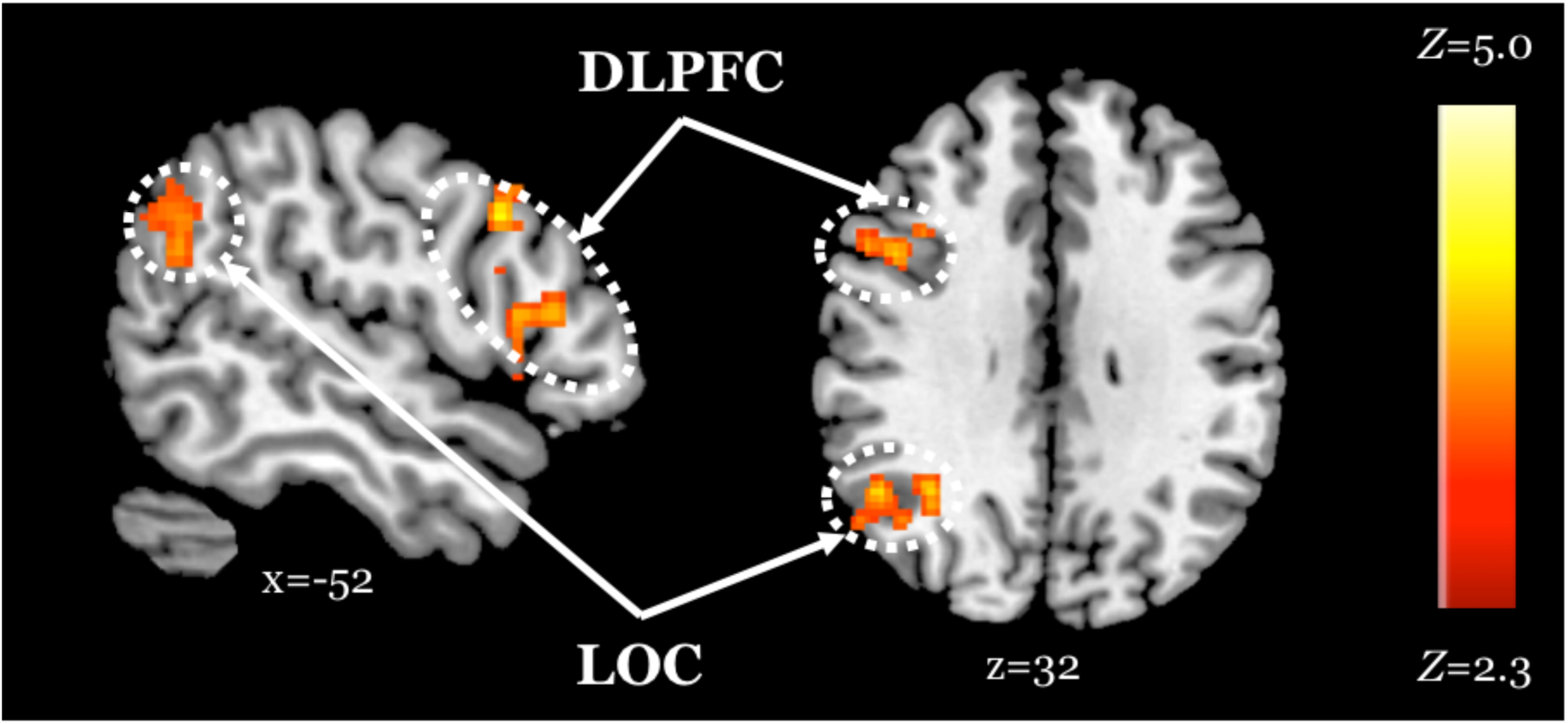
Improved task performance is associated with increased activity within regions of the attentional control network, regardless of age. *Z* statistic maps of regions showing a significant positive association between task performance and Attend Faces > Attend Houses BOLD activity, controlling for age. Images are whole-brain FWE cluster-corrected at *Z* > 2.3 and depicted in MNI space and radiological orientation. DLPFC, dorsolateral prefrontal cortex; LOC, lateral occipital cortex.

**Table 2.** Age group difference local maxima statistics. Hem, hemisphere; B, bilateral; L, left; R, right; x, y, z, coordinates of local maxima in MNI space; *Z*, maximum *Z* statistic; *k*, cluster extent.

| **Contrast** | **Anatomical Region** | **Hem** | **x** | **y** | **z** | **Z** | **k** |
| --- | --- | --- | --- | --- | --- | --- | --- |
| Old<Young | Posterior cingulate cortex | B | -4 | -80 | 8 | 4.91 | 1088 |
|  | Orbitofrontal cortex | L | -34 | 14 | -26 | 4.29 | 878 |
| Old>Young | Fusiform cortex | R | 24 | -42 | -16 | 7.47 | 17997 |
|  | Fusiform cortex | L | -28 | -48 | -10 | 6.93 | 1841 |
|  | Dorsolateral prefrontal cortex | R | 28 | 4 | 54 | 5.24 | 1174 |
| Task Accuracy | Dorsolateral prefrontal cortex | R | 56 | 26 | 22 | 4.06 | 395 |
|  | Lateral occipital cortex | R | 42 | -44 | 22 | 3.85 | 405 |

### Younger but not older adults show a positive association between *OXTR* methylation and neural response during selective social attention

We find a significant interaction between age group and *OXTR* methylation associated with neural response during selective social attention. The slope of the association between *OXTR* methylation and neural response during selective social attention is significantly greater for younger (*n*=98) than older (*n*=85) adults within regions of the attentional control network including anterior cingulate cortex, left dorsolateral prefrontal cortex, and left superior parietal lobule (Figure 4, Table 3). There are no regions in which older adults show a greater slope in the association between *OXTR* methylation and neural response during selective social attention than younger adults.

**Figure 4.**
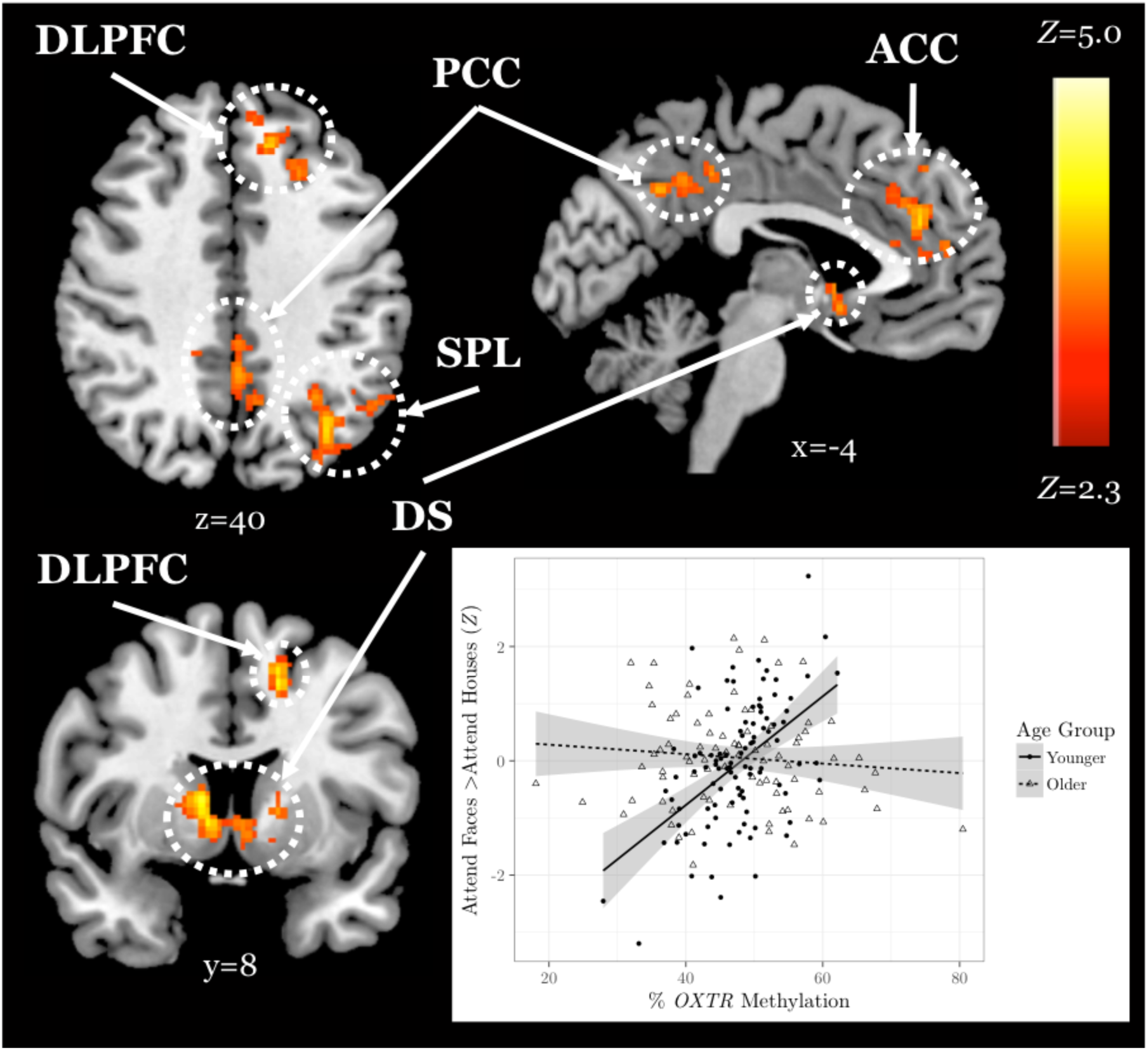
Younger but not older adults show a positive association between *OXTR* methylation and neural response during selective social attention. *Z* statistic maps of regions showing a significant interaction between *OXTR* methylation and age group on Attend Faces > Attend Houses BOLD activity, controlling for task performance. Images are whole-brain FWE cluster-corrected at *Z* > 2.3 and depicted in MNI space and radiological orientation. (Inset) Mean *Z* statistic values are plotted against percent *OXTR* methylation for each participant. Best-fit lines are plotted for younger (solid, n=98) and older (dashed, n=85) adults; grey shading depicts 95% confidence intervals. DLPFC, dorsolateral prefrontal cortex; PCC, posterior cingulate cortex; ACC, anterior cingulate cortex; SPL, superior parietal lobule; DS, dorsal striatum.

**Table 3.** Age group-by-*OXTR* methylation interaction local maxima statistics. Hem, hemisphere; L, left; R, right; x, y, z, coordinates of local maxima in MNI space; *Z*, maximum *Z* statistic; *k*, cluster extent.

| **Anatomical Region** | **Hem** | **x** | **y** | **z** | ***Z*** | ***k*** |
| --- | --- | --- | --- | --- | --- | --- |
| Superior parietal lobule | L | -58 | -68 | 0 | 4.83 | 1270 |
| Dorsolateral prefrontal cortex | L | -18 | 30 | 46 | 4.20 | 472 |
| Dorsal striatum | R | 10 | 6 | 6 | 4.04 | 532 |
| Posterior cingulate cortex | R | 8 | -48 | 36 | 3.67 | 400 |
| Anterior cingulate cortex | L | -2 | 40 | 24 | 3.56 | 533 |

### Social support and *OXTR* methylation interact to account for differences in neural response during selective social attention among older adults

Finally, in an exploratory analysis, we investigate the influence of *OXTR* methylation and social support on neural response during selective social attention within our older adult sample (*n*=84). We find a significant interaction between *OXTR* methylation and reported social support on neural response during selective social attention. To illustrate the interaction, we plot social support by median split and find that older adults who report lower levels of social support display a positive association between *OXTR* methylation and neural activation within right dorsolateral prefrontal cortex, whereas this association is negative for those that report higher levels of social support (Figure 5; MNI coordinates: x=52, y= -18, z=34; *Z*=4.13; cluster extent *k*=1648).

**Figure 5.**
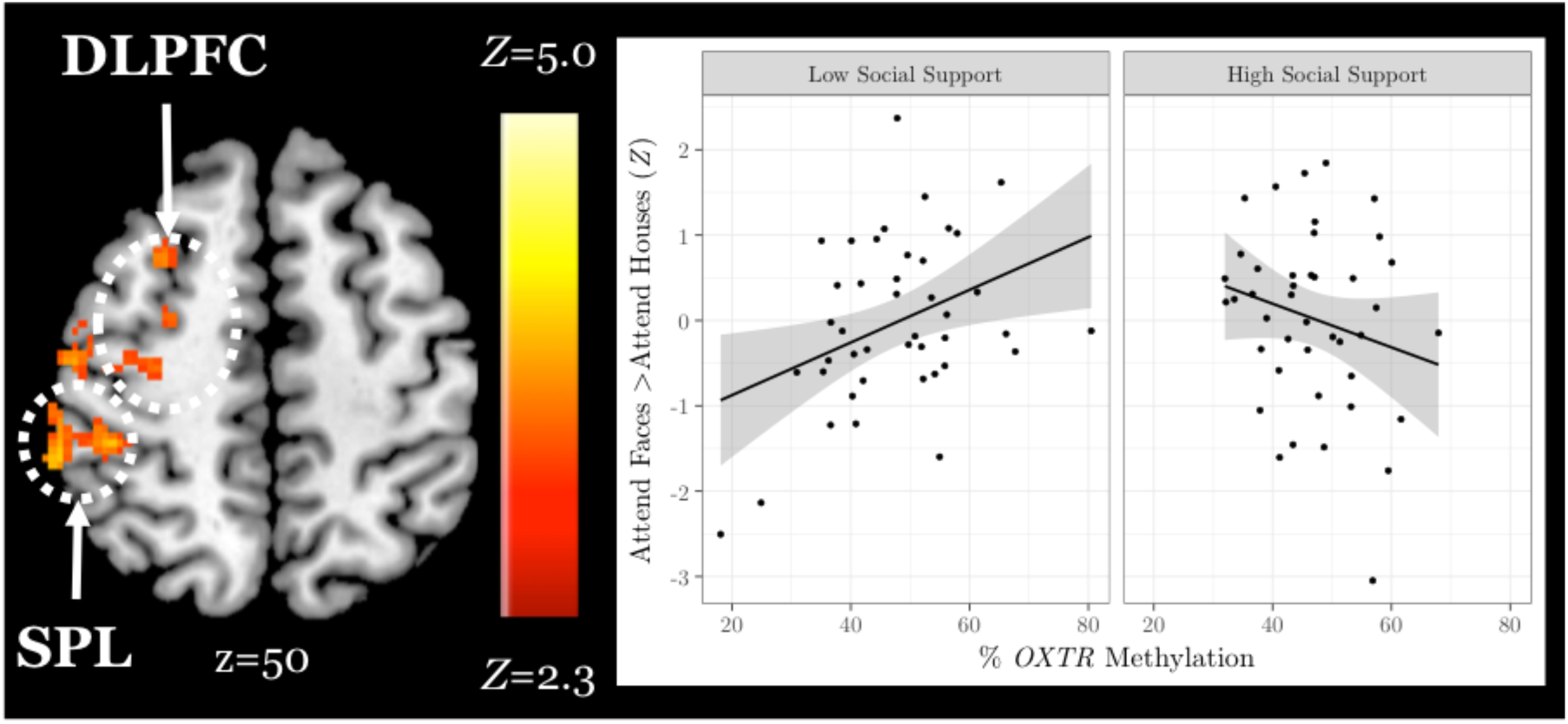
Social support and *OXTR* methylation interact to account for differences in neural response during selective social attention among older adults. *Z* statistic map of regions showing a significant interaction between *OXTR* methylation and self-reported social support on Attend Faces > Attend Houses BOLD activity in older adults (n=84). Images are whole-brain FWE cluster-corrected at *Z* > 2.3 and depicted in MNI space and radiological orientation. (Inset) Mean *Z* statistic values are plotted against percent *OXTR* methylation for each participant. To illustrate the interaction, social support is plotted by median split. Solid lines depict the best-fit line; grey shading depicts 95% confidence intervals. DLPFC, dorsolateral prefrontal cortex; SPL, superior parietal lobule.

## Discussion

In this study, we examine age-related differences in brain response mediated by individual differences in oxytocinergic system function during social attention. After controlling for differences in task performance, we find that older adults activated broad regions of visual and fusiform cortices and recruited dorsolateral prefrontal cortex to a greater extent than younger adults during selective social attention. Older adults also activated orbitofrontal and posterior cingulate cortices, focal regions specialized for social perception (Adolphs, 1999), to a lesser extent than younger adults when attending to faces. These results are consistent with two of the most commonly reported age-related differences in neural activation – the compensatory recruitment of prefrontal regions, and reduced neural specificity, or dedifferentiation, within regions that typically elicit highly specialized responses to specific stimuli in young adults (Park et al., 2004; Reuter-Lorenz and Cappell, 2008; Fabiani, 2012; Sugiura, 2016).

We find that individuals who performed better on the task activated both lateral occipital cortex, a region of the face perception network (Nagy et al., 2012), and dorsolateral prefrontal cortex, a region of the attentional control network (Hopfinger et al., 2000; Seeley et al., 2007), to a greater extent when attending to faces, regardless of age. These results suggest that, across age groups, those who exerted greater attentional “effort” and processed faces to a greater extent performed better on the task.

A whole-brain, age-by-methylation interaction analysis revealed that younger adults with higher levels of *OXTR* methylation, and therefore presumed decreased sensitivity to endogenous oxytocin, recruited regions of the attentional control network to a greater extent when attending to faces. This positive association between *OXTR* methylation and activity within the attentional control network during selective social attention in young adults is robust; here, we replicate an effect previously reported in young adults (Puglia et al., 2018) despite changes in scanner and scanning protocol across our original and replication young adult samples. However, we find no association between *OXTR* methylation and neural response during selective social attention in the older adult sample. This finding is supported by the only other study to date to examine *OXTR* methylation across adulthood; Ebner and colleagues also found associations between site - 934 methylation and social behavior in younger but not older adults (Ebner et al., 2019).

Interestingly, while we find no mean differences in total methylation levels across younger and older adults, methylation levels at site -934 are significantly more variable in our older adult sample – a finding also reported by Ebner and colleagues (Ebner et al., 2019). Early life experiences have been shown to impact *OXTR* methylation at an analogous site in the prairie vole such that pups that received lower levels of parental care within the first few days of life displayed increased levels of *OXTR* methylation in both brain and blood (Perkeybile et al., 2019). It is therefore possible that methylation levels are more widely distributed in the older adult sample because this sample may have a more heterogeneous developmental history.

Alternatively, the aging process itself may result in differences in methylation levels (Richardson, 2003; Koch and Wagner, 2011; Horvath et al., 2012). While we find *OXTR* methylation levels at site -934 are stable in the blood across young adulthood (Connelly and Morris, 2016), interindividual changes have not yet been studied longitudinally into older adulthood. It is important to note that changes in cellular composition associated with aging may confound age-related methylation effects (Jaffe and Irizarry, 2014), although data from our lab suggests that methylation values at site -934 do not vary by cell type (Puglia et al., 2018). Aging is a highly heterogeneous process (Lowsky et al., 2014), and this variability in methylation values in the older adult population could reflect aging differences that are unrelated to chronological age.

We therefore hypothesized that an epigene-by-environment interaction may account for individual differences in neural response during selective social attention in our older adult sample. In an exploratory analysis, we find an interaction between *OXTR* methylation and social support is significantly associated with neural response within dorsolateral prefrontal cortex and superior parietal lobule in older adults. Older adults who reported lower levels of social support resembled the younger adult population; individuals with lower levels of social support and higher levels of *OXTR* methylation exerted greater attentional effort to attend to social information. However, as social support levels increased, the slope between *OXTR* methylation and neural response within attentional control regions decreased. This association provides a potential neurochemical mechanism of behavior in support of the socioemotional selectivity theory. Specifically, older adults tend to be more selective in their relationships, avoiding the formation of new relationships and shedding peripheral social ties to instead invest more in intimate social relationships (Carstensen, 1992; Shaw et al., 2007). Perhaps, older adults who are both highly supported and presumably highly sensitive to endogenous oxytocin (low *OXTR* methylation) continue to exert effort and recruit regions of the attentional control network because they intrinsically find social information worthy of attention and investment of cognitive resources. Conversely, with increasing methylation (presumed less sensitivity to endogenous oxytocin), those who report a supportive social environment may not feel the need to exert extra effort to attend to the social information in this task.

### Limitations and Future Directions

A limitation to the present study is that we did not administer the Social Support Questionnaire to our younger adult samples, so we are unable to determine whether the association we find in our older adult sample is specific to an aging population. Future research directly probing whether age-related shifts in social motivation or specific dimensions of social support (Barrera, 1986) are associated with oxytocinergic system variability and neural function during social perception is warranted.

Our sample had fewer older males (*n*=28) than older females (*n*=60). Therefore, we cannot reliably assess sex differences in the older adult sample, although we find a positive association between *OXTR* methylation and neural response within attentional control regions across sexes in our younger adult sample. The oxytocinergic system, social behavior, and their associated neural systems all show sex differences (Carter et al., 2008); therefore, investigating sex differences in aging will be an important avenue for future research.

Although we controlled for task performance by including this variable as a nuisance regressor in all analyses, future research should set individual threshold levels for task performance so that the neural and epigenetic mechanisms supporting task performance are matched across cognitive demand. However, that task performance varied greatly within both age groups reflects an advantage for our study design as young adults typically score at ceiling on many social cognitive tasks. Finally, to further probe the hypothesized mechanism of *OXTR* methylation and neural response during selective social attention for the theory of socioemotional selectivity, future research should present both familiar and unfamiliar stimuli to understand how age and methylation impact differential activation of attentional control regions during the perception of familiar vs. unfamiliar faces.

### Conclusions

Our results highlight the importance of adopting a lifespan perspective for understanding associations between individual differences in oxytocinergic system function, neural systems supporting social cognition, and social behavior. Maintaining social relationships is critical for longevity, happiness (Holt-Lunstad et al., 2010), and reduced likelihood of developing late-life dementia (Wilson et al., 2007). Particularly as the world’s population continues to age, understanding the neurobiological factors that contribute to healthy social functioning into older adulthood will have critical implications from both a humanitarian and economic standpoint.

### Author Contributions

M.H.P., J.J.C., and J.P.M. designed and conceptualized the study. M.H.P., M.E.L., and M.G.N. wrote the paper. M.H.P. collected the younger adult data and performed the fMRI and epigenetic analyses. M.E.L. collected the older adult data and isolated whole blood DNA for the older adult samples. J.J.C. & J.P.M. provided edits and comments to the manuscript and funding for the work.

## Acknowledgements

We thank Timothy A. Salthouse for providing access to the Virginia Cognitive Aging Project (funded by National Institute on Aging Grant AG024270), and for his helpful edits and comments on the manuscript. We also thank Andrew Graves, Hazel Lindal, Amalia McDonald, Jessica Pham, Tyler Santander, and Brenda Straka for assistance with data collection. This research was supported by National Science Foundation Grants 1228522 and 1657726 to J.J.C. and J.P.M.

